# IQM-22110 as a selective K_V_4.3/KChIP3 modulator. Molecular determinants of the KChIP3 binding site

**DOI:** 10.1101/2025.05.29.656781

**Authors:** Paula G. Socuellamos, Carmen Viedma-Barba, Angela de Benito-Bueno, Maria Angeles Bonache, Maria Ropero, Sara Diez, Pablo Elizalde, Irene Marín-Olivero, Maria Redondo, Jose Ramon Naranjo, Juan A. Gonzalez-Vera, Angel Orte, Angel Perez-Lara, Mercedes Martin-Martinez, Carmen Valenzuela, Marta Gutierrez-Rodriguez

## Abstract

The goal of the present study was to discover novel KChIP ligands as research tools for modulating the K_V_4.3/KChIP channels. By employing a multidisciplinary approach, combining medicinal chemistry and electrophysiology studies, a novel K_V_4.3/KChIP modulator (IQM-22110) was successfully identified. IQM-22110 has emerged from the combination of our prior knowledge regarding the (phenylacetamido)benzoic acid moiety as an effective scaffold for KChIP3 ligands and a virtual screening of a focused chemical library. Guided by docking studies—which indicated that incorporating an additional aromatic ring could enhance binding affinity—IQM-22110 was selected for synthesis and identified as a potent KChIP3 ligand. Its electrophysiological effects on K_V_4.3/KChIP3 currents indicate that IQM-22110 binds to a high affinity site in K_V_4.3/KChIP3 channels that it is not present in K_V_4.3/KChIP2 or K_V_4.3. To the best of our knowledge, here we describe the first KChIP3 ligand that selectively modulates K_V_4.3/KChIP3 *versus* K_V_4.3/KChIP2 and K_V_4.3 alone channels. Given that KChIP2 is primarily expressed in heart, our findings might pave the way for the development of K_V_4.3/KChIP3 blockers with reduced cardiac side effects. Computational and site-directed mutagenesis studies allowed the identification of IQM-22110’s binding site on KChIP3. Knowledge gained from our structural and functional studies with this novel KChIP3 ligand could establish the basis for drug discovery programs fostering treatments for diseases in which K_V_4.3/KChIPs channels are involved.

## 1. Introduction

Channelopathies arising from alterations in voltage-dependent potassium K_V_4 currents have been implicated in a range of pathological conditions, including Brugada syndrome, cardiac hypertrophy, atrial fibrillation, heart failure, schizophrenia, epilepsy or dementia [1–3].

K_V_4 channels are the main contributors to the cardiac transient outward K^+^ current (*I*_tof_), and the A-type potassium currents (*I*_A_) in nerve cells. Consequently, K_V_4 channels play a key role in the early phase of repolarization in cardiac action potentials (AP) [4] and, in the nervous system, they regulate the action potential waveform and duration, latency to the first spike, and the timing of synaptic inputs [5]. K_V_4 channels comprised three members (K_V_4.1, K_V_4.2 and K_V_4.3), that to fully reproduce the *I*_tof_ and *I*_A_ currents, need to assemble with other accessory subunits [6], including KChIPs (potassium channel interacting proteins), DPPs (dipeptidyl-peptidase-like proteins), KCNEs (potassium voltage-gated channel subfamily E regulatory subunits), KChAPs and Kvβx accessory subunits, forming channelosomes [7–9].

There is an ongoing effort to understand the role of the K_V_4 channelosomes in human pathophysiology and the pharmacological consequences of its modulation by small molecules [2,3,10–16].

KChIPs are members of the calcium-binding protein superfamily. In mammals, the KChIP subfamily consists of four members: KChIP1, KChIP2, KChIP3 (also known as DREAM or calsenilin), and KChIP4 (also known as calsenilin-like protein). These proteins are encoded by four genes (*KCNIP1-4*), each with multiple spliced variants. While KChIP1-4 are found in the brain, KChIP2 is the most prominent in the heart. KChIP1-3 play a crucial role in enhancing the trafficking of K_V_4.3 channels to the plasma membrane, delaying their inactivation kinetics, and accelerating both activation and recovery from inactivation kinetics [2], while KChIP4 hinders K_V_4 trafficking to the plasma membrane [17]. To gain deeper insights into the structural basis of K_V_4 recognition and modulation by KChIPs, 3D structures of several KChIP-K_V_4 complexes have been resolved [18–22].These structural insights, combined with the available three-dimensional (3D) structures of KChIPs in their unbound state (PDB IDs: 1S1E, 2E6W, 2JUL, 3DD4) provide a valuable framework for the identification of novel KChIP ligands and their potential binding sites [23–26].

The study of the K_V_4 channelosome and its regulation by other proteins through novel and selective small molecule ligands presents both pharmacological and molecular challenges. In this regard, considering that KChIP ligands significantly influence the overall biophysical properties of K_V_4.3/KChIP currents, the identification of small molecules targeting KChIPs and modulating the K_V_4.3 electrophysiology have garnered significant interest [2,3,10,27–29].

The objective of this study was to discover novel KChIP ligands as research tools for modulating the K_V_4.3 channelosome. By employing a multidisciplinary approach, combining medicinal chemistry and electrophysiology studies, a novel K_V_4.3/KChIP modulator (IQM-22110) was successfully identified. IQM-22110 has emerged from the combination of our prior knowledge regarding the (phenylacetamido)benzoic acid moiety as an effective scaffold for KChIP3 ligands and a virtual screening of a focused chemical library. Guided by docking studies—which indicated that incorporating an additional aromatic ring could enhance binding affinity—IQM-22110 was selected for synthesis and identified as a potent KChIP3 ligand. Its electrophysiological effects on K_V_4.3/KChIP3 channels indicate that IQM-22110 binds to a high affinity site in K_V_4.3/KChIP3 that it is not present in K_V_4.3/KChIP2. Given that KChIP2 is primarily expressed in heart, our finding paves the way for reducing cardiac side effects through the selective modulation of the K_V_4.3 current mediated by KChIP3. Computational and site-directed mutagenesis studies allowed the identification of IQM-22110’s binding site on KChIP3. Knowledge gained from our structural and functional studies with this novel KChIP3 ligand could establish the basis for drug discovery programs fostering treatments for diseases that involve K_V_4-mediated-channelopathies.

## 2. Results

### Medicinal chemistry: Design and synthesis of novel KChIP ligands

Our previous structure-activity relationship studies identified a (phenylacetamido)benzoic acid moiety, as suitable skeleton for KChIP3 ligands, where the introduction of additional substituents in the benzoic acid ring provides the necessary features for an efficient interaction with KChIP3 [27]. Inspired by this knowledge, we have designed a virtual focused chemical library of 26 novel compounds with the aim to incorporate substituents on the phenylacetamido moiety (R_1_ and R_2_, Figure 1A) potentially capable to bind unexplored cavities on KChIP3 (Figure 1B). The virtual library was filtered by molecular docking studies with several of our reported homology models of human *h*KChIP3 [27], using Glide and the induce fit docking (IFD) protocol within the Schrödinger Suite of programs [30,31], that allows the flexibility of the ligand and the receptor amino acid side chains in a window of 5 Å from the ligand. Based on our previous studies on the KChIP3-ligand’s binding site, we selected a hydrophobic pocket around Tyr118 and Tyr130 residues for the molecular docking studies. These residues, located approximately at the center of a large hydrophobic cleft in KChIP3, were essential for the binding of our earlier ligands.

**Figure 1.**
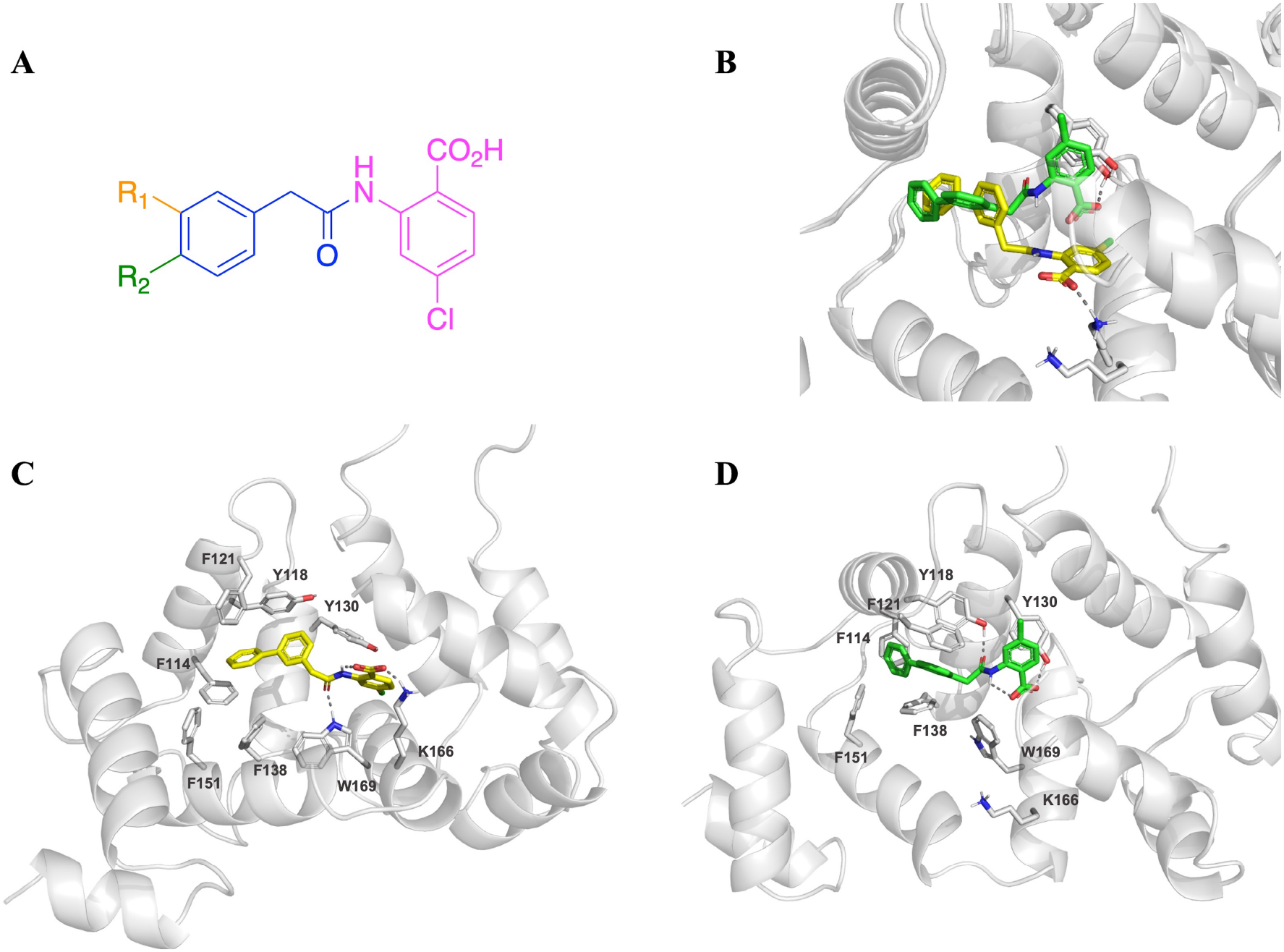
Selection of potential KChIP3 ligands guided by IFD studies. **A**. General structure of the virtual chemical library compounds. **B**. Superimposition of the two representatives IFD poses. **C**. KChIP3 – ligand complex pose A, showing the hydrogen bond with Trp169. **D**. KChIP3 – ligand complex poses B,displaying the hydrogen bond with Tyr118. Hydrogen bonds are depicted as grey dotted lines. For clarity, only polar hydrogens are shown

The analysis of the IFD results showed that, in general, the designed compounds adopt two different binding modes, mainly differing in the pocket that accommodates the benzoic acid moiety. In one pocket, a hydrogen bond is stablished between the carboxylic acid and Tyr130 (pose A), while in the other a salt bridge is formed with Lys166 (pose B) (Figure 1 C, D). Notably, despite these variations in the benzoic acid positioning, derivatives containing a 2-biphenyl-acetamide moiety consistently localize the biphenyl group within the same hydrophobic cavity surrounded by Phe114, Tyr118, Phe121, Phe138 and Phe151. For this biphenyl derivative, a hydrogen bond is observed between the carbonyl group of the central amide linker and either Trp169 (in pose A) or Tyr118 (in pose B) (Figure 1B-C). Overall, these results highlight the potential benefits of incorporating a phenyl ring at R_1_ position, as it leads to additional interactions within the hydrophobic cavity, which may enhance compound affinity for KChIP3 (Figure 1). Based on this premise, we decided to explore the impact of the introduction of a phenyl ring at R_1_ on the KChIP3 binding and on the modulation of the K_V_4.3/KChIP3 current.

To this end, IQM-22110 (2-(2-([1,1’-biphenyl]-3-yl)acetamido)-4-chlorobenzoic acid) was synthesized using a convergent strategy, involving the reaction between methyl 2-amino-4-chlorobenzoate and biphenyl carboxylic acid **3** followed by hydrolysis of the corresponding methyl ester (Figure 2A). Biphenyl carboxylic acid **3** was obtained via a Suzuki cross-coupling reaction between 2-(3-bromophenyl)acetic acid (**1**) and the corresponding boronic acid **2**.

**Figure 2.**
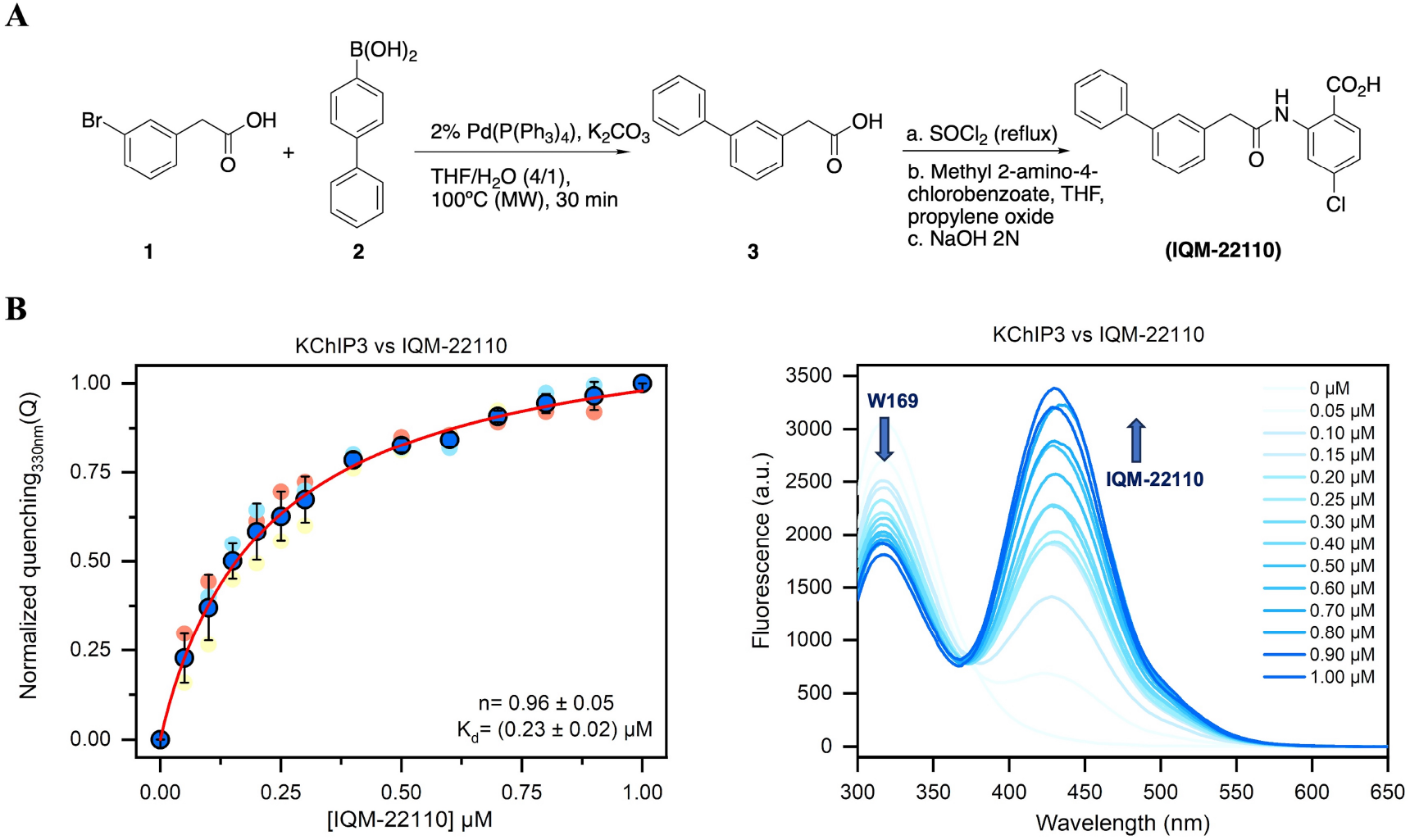
Synthesis of IQM-22110 and evidence of its binding to KChIP3. **A**. Synthesis of IQM-22110. **B**. Concentration-dependent interaction between IQM-22110 and KChIP3. Left panel: Plot of the tryptophan quenching by FRET towards IQM-22110. Data from the three repetitions (open symbols) are shown as well as the average value (black-filled symbols) and error bars as s.d. The red line represents the fit to a Hill equation. Right panel shows the fluorescence emission spectra (λex = 260 nm) from a representative titration of KChIP3 (0.5 μM) with increasing concentrations of IQM-22110.

Taking advantage of the fact that KChIP proteins possess a single tryptophan residue within the ligand-binding hydrophobic cleft, the binding of IQM-22110 to KChIP3 was assessed by monitoring the quenching of intrinsic tryptophan fluorescence. As it is shown in Figure 2A, the occurrence of fluorescence resonance energy transfer (FRET) is evidenced by a decrease in tryptophan fluorescence intensity accompanied by a concomitant increase in the emission of IQM-22110, indicating close spatial proximity between the ligand’s chromophore and the Trp169 of KChIP3.

To assess the affinity of IQM-22110 by KChIP3, we plotted the quenching of tryptophan fluorescence emission at 330 nm as a function of compound concentration (Figure 2B). The resulting data were fitted using the Hill equation (see Methodology), yielding a dissociation constant (K_d_) of 230 ± 20 nM and a Hill coefficient close to 1, indicating a non-cooperative binding to KChIP3 and supporting one single binding site, at least at FRET-allowing distances. Based upon these results, IQM-22110 represents a novel pharmacological tool to study the modulation of KChIP3 and explore its therapeutic relevance.

Using the spectral properties of KChIP3 tryptophan fluorescence and the absorption characteristics of IQM-22110, we calculated a Förster distance (R_0_) of 10.55 ± 0.09 Å (assuming free rotation of the dyes, which is likely not a totally valid assumption but provides a rough estimation of the distance). At saturating concentrations of compound, the extent of tryptophan fluorescence quenching was estimated to be around 0.58 ± 0.17. This FRET efficiency corresponded to an apparent distance tryptophan–ligand of circa. 10.02 ± 0.78 Å

### Concentration-dependent inhibition of IQM-22110 on K_V_4.3, K_V_4.3+KChIP3 and K_V_4.3+KChIP2 currents

To explore the possible electrophysiological effects of IQM-22110 on Kv4-KChIPx channels, we analyzed its potential blocking effects on K_V_4.3 and K_V_4.3/KChIP3 channels. To assess this issue, we applied test pulses from -80 mV to +60 mV and we measured the peak current and the charge (measured as the area under the current) in the absence and in the presence of different IQM-22110 concentrations (between 0.001 and 150 μM) (Figure 3).

**Figure 3.**
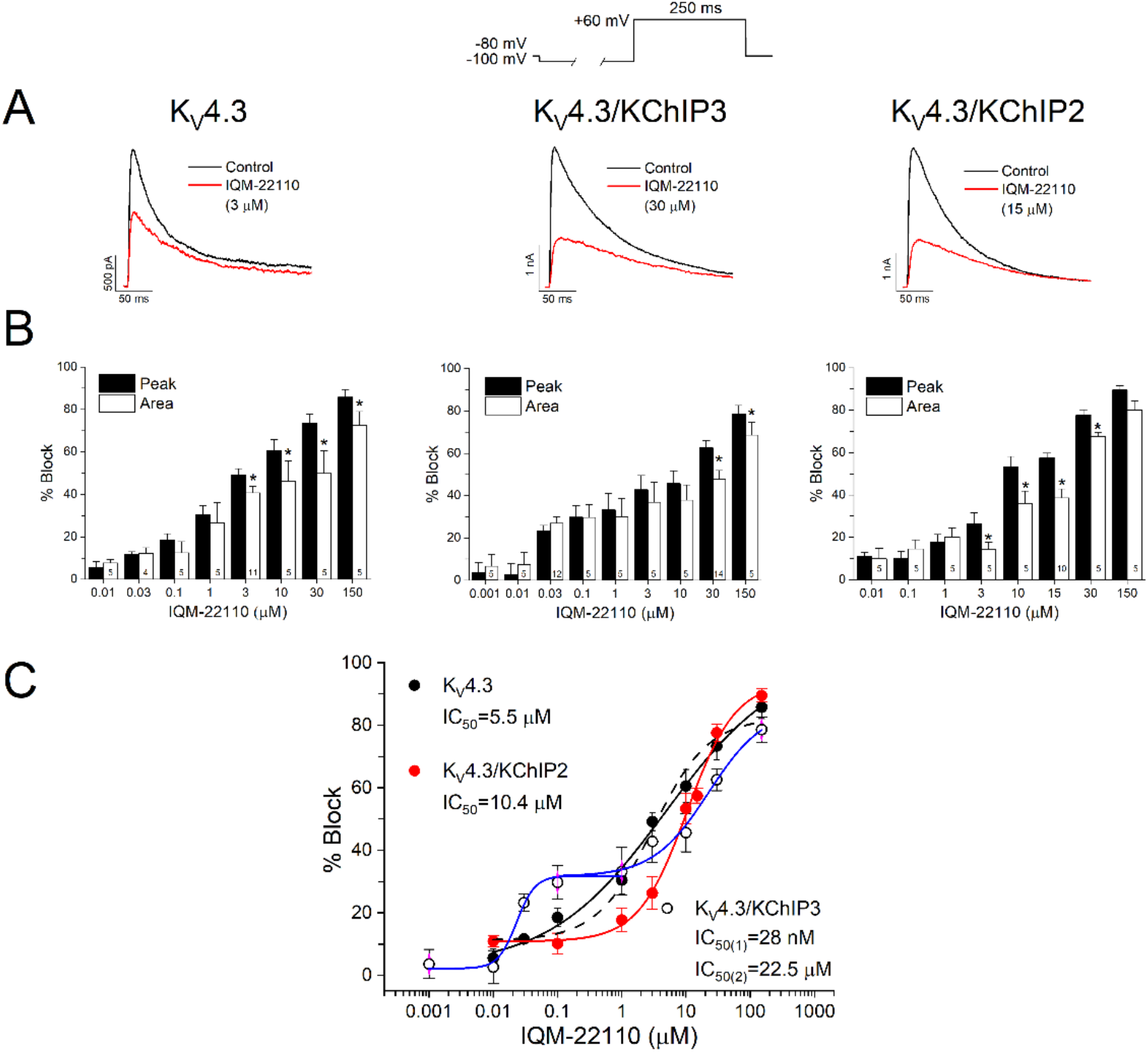
Concentration-dependent block produced by IQM-22110 on K_V_4.3, K_V_4.3/KChIP3 and K_V_4.3/KChIP2 channels. **A**. Current traces in the absence and in the presence of IQM-22110 are shown. **B**. Effects of IQM-22110 (between 0.001 and 150 μM) on K_V_4.3, K_V_4.3/KChIP3 and K_V_4.3/KChIP2 currents measured at the maximum peak current and on the charge (measured as the area under the current) during the application of a 250 ms pulse to +60 mV (n = 4 – 12, \**P* < 0.05 when comparing the effect of IQM-22110 on the peak current and on the charge through K_V_4.3, K_V_4.3/KChIP3 and K_V_4.3/KChIP2 channels). **C**. Concentration-block of K_V_4.3, K_V_4.3/KChIP3 and K_V_4.3/KChIP2 channels induced by IQM-22110 (n = 4 – 12).

Figure 3A shows original current traces obtained after the activation of K_V_4.3, K_V_4.3/KChIP3 and K_V_4.3/KChIP2 channels in the absence and in the presence of equipotent IQM-22110 concentrations (3, 30 and 15 μM, respectively). Under these experimental conditions, this compound decreased the peak current and the charge during the application of 250 ms pulses to +60 mV. Also, IQM-22110 slowed the inactivation process (Figure 3B).

As it can be observed, IQM-22110 inhibited the K_V_4.3 current with a IC_50_ of 5.5 μM measured as the decrease in the maximum peak current. However, the concentration-dependent block of K_V_4.3/KChIP3 channels induced by this compound was found to be biphasic with two IC_50_ values at 0.03 and 22.5 μM, suggesting that this new ligand exhibits very high selectivity for KChIP3 at low concentrations. To corroborate this finding and further explore its KChIP3 selectivity, we similarly investigated the concentration-dependent block of IQM-22110 on K_V_4.3/KChIP2 channels, which was found to be monophasic with a IC_50_ value of 10.4 μM (Figure 3C). This finding highlights the high selectivity of IQM-22110 towards K_V_4.3/KChIP3 channels compared to K_V_4.3 or K_V_4.3/KChIP2 channels.

### Voltage-dependence of IQM-22110 on Kv4.3 and Kv4.3/KChIP3 channels

In order to analyze the voltage-dependent of block induced by IQM-22110 on K_V_4.3 and K_V_4.3/KChIP3 channels, the pulse protocol shown in the upper panel of Figure 4 was used. Cells were held at -80 mV and 250 ms depolarizing pulses from -80 to +60 in 10 mV steps every 10 s were applied. The current amplitude was measured at the maximum peak current and these values were plotted versus the test pulse voltage. Figure 4A shows superimposed potassium current traces through K_V_4.3 in the absence and in the presence of IQM-22110 (3 μM). Figure 4B shows K_V_4.3/KChIP3 current traces in the absence and in the presence of IQM-22110 concentrations close to the IC_50(1)_ and IC_50(2)_ values exhibited by (30 nM and 30 μM).

**Figure 4.**
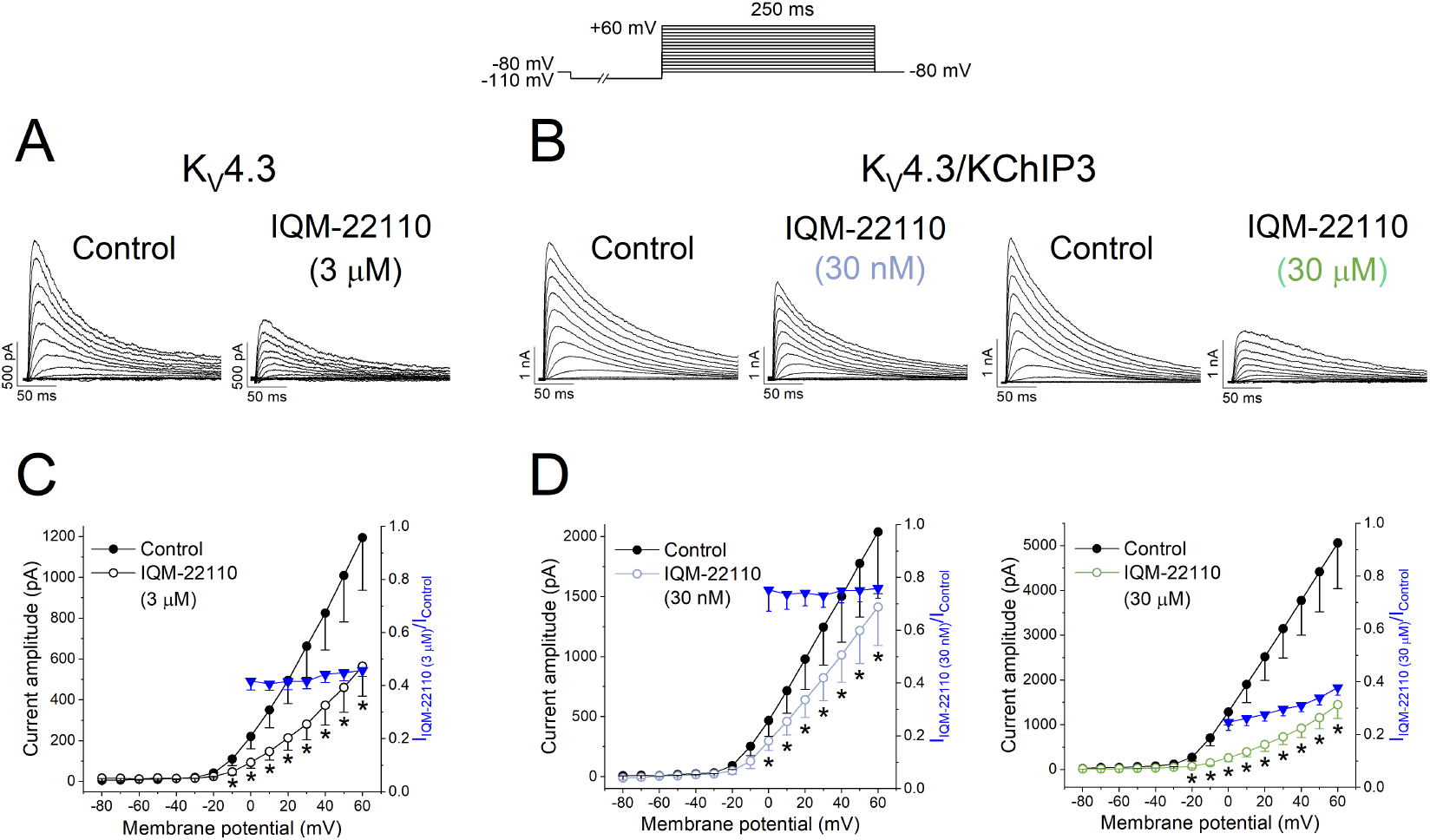
Voltage-dependent effects of IQM-22110 on K_V_4.3 and K_V_4.3/KChIP3 channels. **A**. Representative recordings obtained in the absence and in the presence of IQM-22110 in K_V_4.3 (3 μM), andK_V_4.3/KChIP3 (30 nM and 30 μM) channels after applying the pulse protocol shown at the top of the figure. **C**. I-V relationship of the currents generated by K_V_4.3 in the absence and in the presence of IQM-22110 (3 μM) and **D**. by K_V_4.3/KChIP3 channels in the absence and in the presence of IQM-22110 at 30 nM and 30 μM respectively when measured at the maximum peak current during the application of the depolarizing pulse protocol shown in the upper part of the figure. The right y-axis of both graphs shows the relative current obtained after dividing the current magnitude obtained in the presence of IQM-22110 by that measured in the absence of the compound. (n = 5 – 6, \**P* < 0.05 when comparing data in presence of IQM-22110 with control).

Under these experimental conditions, IQM-22110 (3 μM and 30 μM) decreased the amplitude of the current at membrane potentials positive to -10 mV (in K_V_4.3 and K_V_4.3/KChIP3) slowing the inactivation kinetics. However, IQM-22110 (30 nM) on K_V_4.3/KChIP3 channels slightly accelerated the inactivation kinetics (72.1 ± 6.1 ms *versus* 64.8 ± 4.5 ms in the absence and in the presence of IQM-22110, n = 5, *P* > 0.05).

Block induced by IQM-22110, in the absence of KChIP3, was voltage-independent, arising similar degree between 0 and +60 mV. Similarly, IQM-22110 (30 nM) produced a voltage-independent block of K_V_4.3/KChIP3 channels. However, IQM-22110 (30 μM) block was maximum at 0 mV and significantly decreased at positive potentials (blue symbols in Figure 4, n = 5, *P* < 0.05), when the membrane potential was more positive and, thus, the likelihood of opening is higher. These results suggest that the block induced by nM concentrations (IC_50(1)_) of IQM-22110 on K_V_4.3/KChIP3 is consistent with the block of a closed-activated state, whereas that on K_V_4.3/KChIP3 channels is consistent with an open/closed or inactivated state block, whereas that observed by IQM-22110 in the μM range (IC_50(2)_) on K_V_4.3/KChIP3 is consistent with the block of a closed-activated state.

### Use-dependent block and recovery-dependent of inactivation

In order to assess the use-dependent inhibition produced by IQM-22110 on K_V_4.3 and K_V_4.3/KChIP3, K_V_4.3/KChIP3, train protocols, consisting of 15 pulses from -80 mV to +50 mV of 25, 250 or 500 ms at a frequency of 2 Hz were applied (Figure 5).

**Figure 5.**
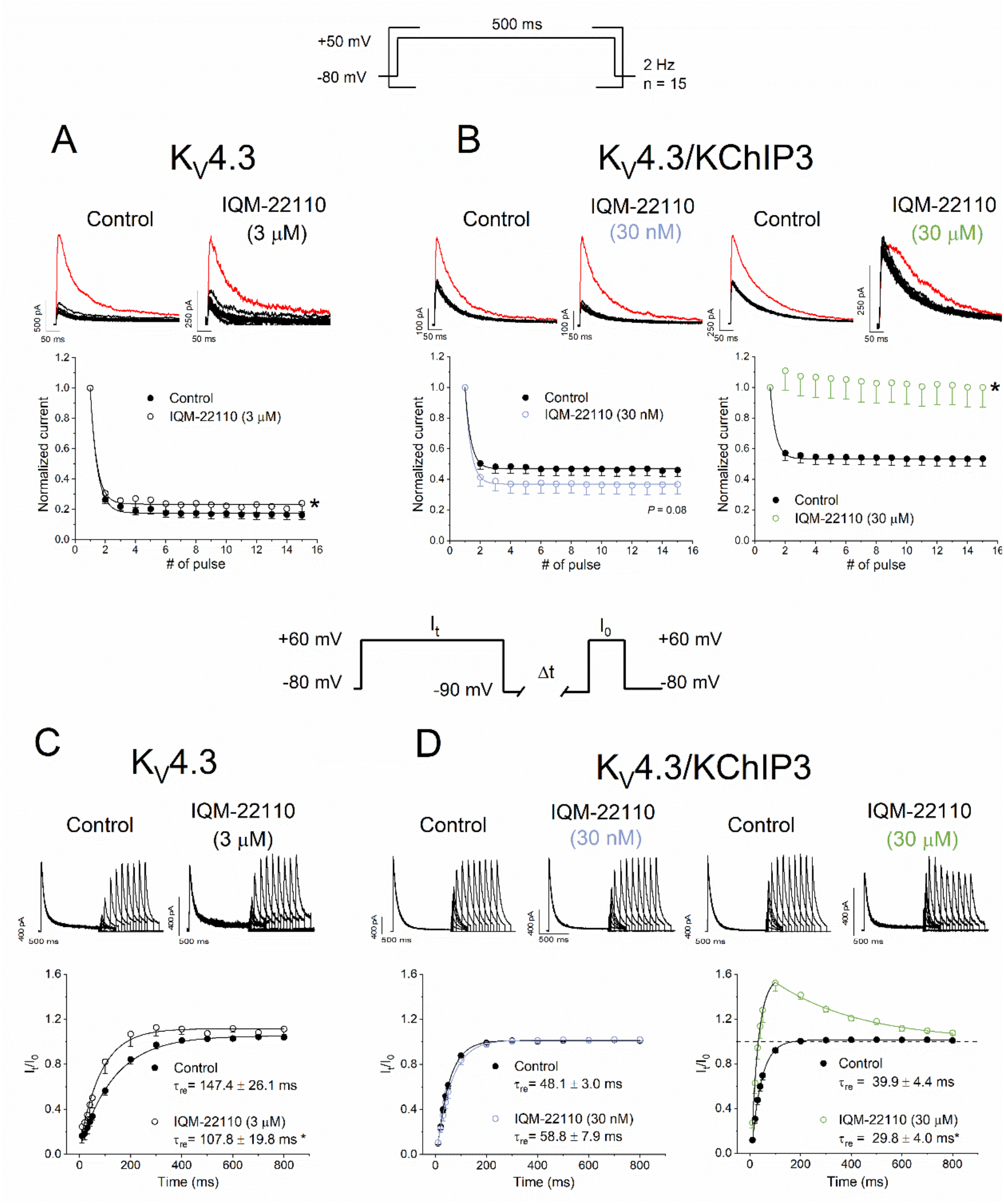
Use-dependent and recovery from inactivation of block induced by IQM-22110 on K_V_4.3 and K_V_4.3/KChIP3 channels. **A**. Original recordings and plot of the normalized values obtained during train application in the absence and in the presence of IQM-22110 in K_V_4.3, and **B**. K_V_4.3/KChIP3 channels after applying a pulse-train protocol (500 ms, 2 Hz) in the absence and in the presence of IQM-22110 at 30 nM (blue) and at 30 μM (green) shown in the top panel. In both cases, the first pulse of each recording is shown in red. (n = 5 – 6. *: *P* < 0.05 when comparing the degree of decay of the current in presence of IQM-22110 vs. control conditions). **C**. Representative recordings obtained after applying the pulse protocol shown in the figure in the absence and in the presence of IQM-22110 at 3 μM in K_V_4.3 and **D**. 30 nM (blue) and 30 μM (green) in K_V_4.3/KChIP3 channels. Data obtained after plotting the I_t_/I_0_ vs. the interpulse period separating both the conditioning and the test pulse is shown in the lower part of the figure. Data were fitted to an exponential fit in order to calculate the time constant of recovery (τ_re_) (n = 5 – 6. * *P* < 0.05 when comparing data in presence of IQM-22110 with control).

In the absence of KChIP3, trains of 500 ms-pulses at 2 Hz induced a huge decrease under control conditions and IQM-22110 (3 μM) decreased this exponential decay from 83.8 ± 2.8% to 75.0 ± 5.1%, n = 5, *P* < 0.05) (Figure 5A). When KChIP3 was present, the current decay during train application under control conditions was lower than that observed in the absence of KChIP3, due to the faster recovery from inactivation induced by this regulatory subunit (Figure 5B). IQM-22110 (30 nM) slightly increased the use-dependent decay during the train pulse application (63.3 ± 6.4% *versus* 53.9 ± 4.1% in the absence and in the presence of IQM-22110, n = 5, *P* = 0.08). On the other hand, IQM-22110 (30 μM) abrogated the use-dependent decay of the K_V_4.3/KChIP3 current observed under control conditions. The results obtained in K_V_4.3 and in K_V_4.3/KChIP3 with 3 and 30 μM, respectively, may be explained by an acceleration of the recovery of inactivation; whereas the slightly increase in the percentage of the current decay for K_V_4.3/KChIP3 in the presence of IQM-22110 (30 nM) during the train application could be due to a slower recovery from inactivation.

To assess this issue, we analyzed the recovery process of inactivation in K_V_4.3 and in K_V_4.3/KChIP3 currents in the absence and in the presence of IQM-22110 (3 μM for K_V_4.3, and 30 nM or 30 μM for K_V_4.3/KChIP3 channels) (Figure 5) by applying the double-pulse protocol shown in Figure 5C. As it is observed, the time constant of recovery of inactivation (τ_re_) of K_V_4.3 was accelerated by IQM-22110 both in the absence and in the presence of KChIP3 (*P* < 0.05) in the presence of 3 and 30 μM, respectively, and induced an overshoot both in K_V_4.3 and in K_V_4.3/KChIP3 channels in the presence of IQM-22110 at 3 μM or 30 μM, respectively. These results suggest that IQM-22110 binds preferentially to a closed or closed/activated state, with low or null affinity for the inactivated state. Thus, at the end of a 1 s depolarizing pulse (that promotes the inactivation of most K_V_4.3 channels with or without KChIP3), the compound dissociates from the channel and, depending on its biophysical properties, an overshoot more or less pronounced will appear, likely representing the dissociation of IQM-22110 from the inactivated state of the channel. However, the recovery process of inactivation in the presence of IQM-22110 (30 nM) was slightly slowed (58.8 ± 7.9 ms *versus* 48.1 ± 3.0 ms, n = 5, *P* = 0.08), suggesting that, at this concentration, IQM-22110 is binding to another structure different from the K_V_4.3 channel, likely KChIP3.

### Deciphering the binding site of IQM-22110 in KChIP3

IQM-22110 has proven to be a suitable ligand for KChIP3 presenting a novel and high affinity site in their binding to K_V_4.3/KChIP3. Therefore, we set out to study its binding mode in more detail and correlate computational studies with site-directed mutagenesis studies by using electrophysiological techniques. Given that IFD studies suggested Tyr130 and Lys166 as key residues, and that a hydrogen bond was observed between the carbonyl group of the central amide linker of IQM-22110 and either Trp169 (in pose A, Figure 1C) or Tyr118 (in pose B, Figure 1D), we selected three KChIP3 mutants to study the blocking effects of IQM-22110 at 30 nM on K_V_4.3, K_V_4.3/KChIP3^WT^, K_V_4.3/KChIP3^Y118A^, K_V_4.3/KChIP3^Y130A^ and K_V_4.3/KChIP3^K166A^.

As is shown in Figure 6, IQM-22110 at 30 nM inhibited the K_V_4.3/KChIP3^WT^ by 23.2 ± 2.7% (n = 12). On the other hand, this ligand produced a decrease of 9.7 ± 2.5 (n = 5, *P* < 0.05) and 9.0 ± 2.5% (n = 5, *P* < 0.05) in the K_V_4.3/KChIP3^Y130A^ and K_V_4.3/KChIP3^K166A^, respectively. However, the inhibition produced by IQM-22110 on K_V_4.3/KChIP3^Y118A^ current was similar to that of K_V_4.3/KChIP3^WT^ (17.0 ± 3.6%, n = 5, *P* > 0.05). At 30 μM, IQM-22110 induced the same degree of blockade in K_V_4.3/KChIP3^WT^ channels as in K_V_4.3/KChIP3^Y118A^, K_V_4.3/KChIP3^Y130A^ or K_V_4.3/KChIP3^K166A^ channels, indicating that these three residues indeed represent the molecular determinants of the high-affinity binding site. All these results support the pose A as the most likely binding mode (Figure 1C).

**Figure 6.**
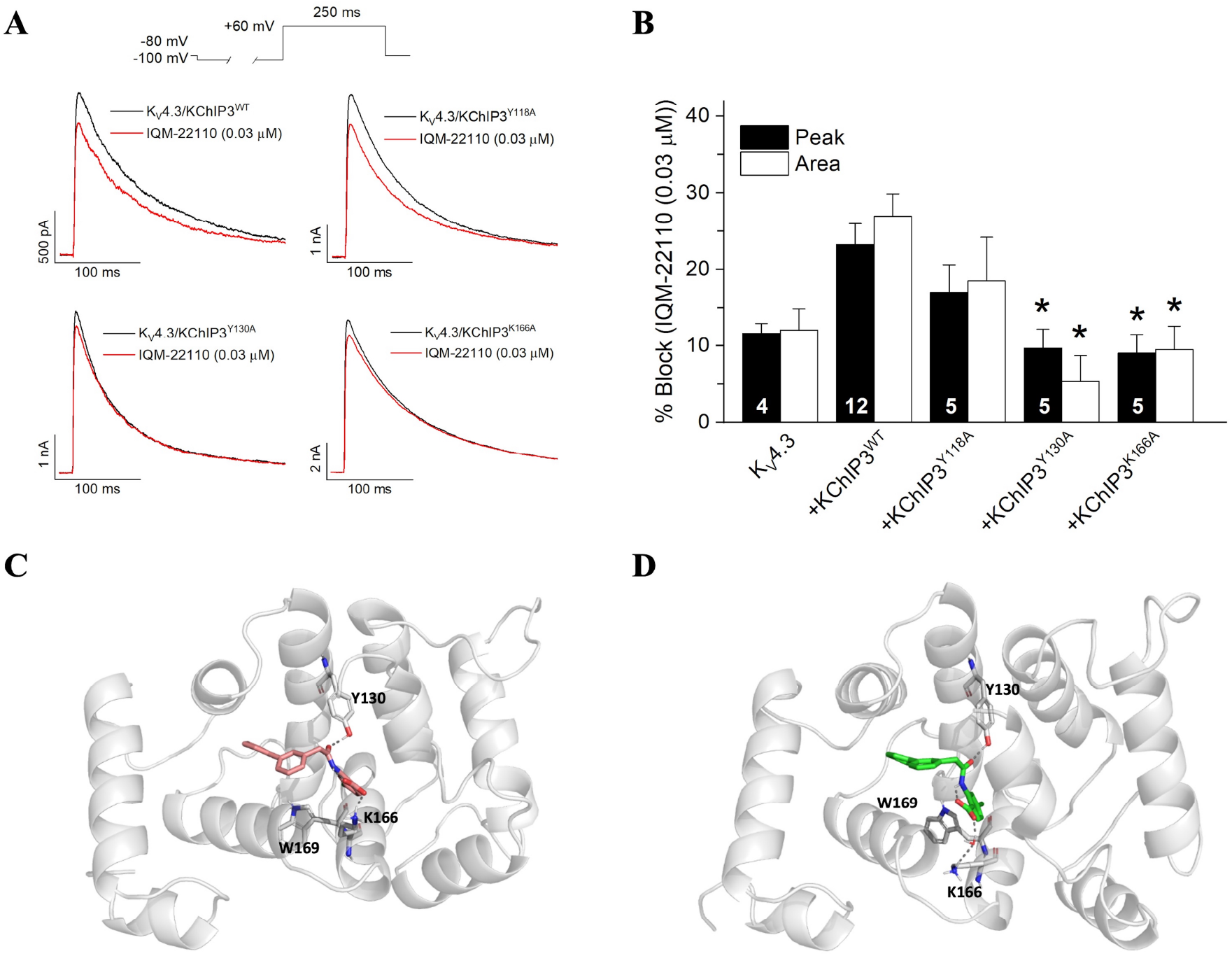
IQM-22110 binding site. **A**. Blocking effects of IQM-22110 (0.03 μM) on K_V_4.3, K_V_4.3/KChIP3^WT^, K_V_4.3/KChIP3^Y118A^, K_V_4.3/KChIP3^Y130A^ and K_V_4.3/KChIP3K^166A^. Original records in the absence and in the presence of IQM-22110. **B**. Bar-graph representing the degree of block in each experimental condition. (n = 4 – 12. \**P*< 0.05 when comparing with block produced in K_V_4.3/KChIP3^WT^).Representative KChIP3-IQM-22110 1000 ns frame with a salt bridge between Lys166 and the ligand carboxylic oxygen. **D**. Representative KChIP3-IQM22110 1000 ns frame with a hydrogen bond with Lys166 mediated by a water molecule.

To better delineate the binding site, starting from pose A (Figure 1C) in which Tyr118 is not involved in the ligand binding, we performed molecular dynamic (MD) studies which allows the flexibility of both the ligand and the whole protein. Toward this end the KChIP3-IQM-22110 complex was solvated in an octahedral box of TIP3P water molecules and neutralized. Three 1000 ns MD replicas were performed to ensure the reproducibility of the results. Then, we analyzed salt bridges and hydrogen bonds as they are relevant interactions for stabilizing the complex. MDs reveal that within the first 100 ns, the amide carbonyl group of the ligand swifts away from Trp169, getting close to Tyr130 and stablishing a new hydrogen bond with this residue (Figure 6). This interaction, absent in the docking pose, is in agreement with mutagenesis studies and FRET analysis. This hydrogen bond is maintained in over 78 % of the 1000 ns simulation. Additionally, the COOH oxygen of IQM-22110 forms a salt bridge with Lys166 side chain in over 57% of the MDs. Interestingly, when a direct interaction is absent, a water-mediated hydrogen bond was present in at least 13% of the MD. Considering both direct and water-mediated interactions, IQM-22110 is interacting with Lys166 in over 70% of the simulation. Additionally, the biphenyl moiety is within a hydrophobic pocket, that comprised several aromatic residues Phe114, Tyr118, Phe121, Phe122, Tyr130, Phe138 and Phe151. All these results suggest that Tyr130 and Lys166 in KChIP3 are critical residues for the binding of IQM-22110 to the high affinity site in K_V_4.3/KChIP3.

## 3. Discussion

In the present study, we have designed and synthesized for the first time a KChIP3 ligand, IQM-22110, that selectively modulates K_V_4.3/KChIP3 *versus* K_V_4.3/KChIP2 and K_V_4.3 alone channels. We demonstrated that IQM-22110: (a) binds to K_V_4.3/KChIP3 channels in a concentration-, time- and voltage-dependent manner; (b) inhibits the maximum peak current during depolarization; (c) at low concentrations (nM range), IQM-22110 blocks K_V_4.3/KChIP3, but not K_V_4.3 nor K_V_4.3/KChIP2, suggesting that in this range of concentrations, this ligand is selective for KChIP3; and finally (d) the molecular determinants of the KChIP3 binding site have been identified.

K_V_4.3 is a voltage-gated potassium channel subunit essential for generating transient outward potassium current in the heart (*I*_tof_) and in the brain (*I*_A_). In cardiac tissue, K_V_4.3 contributes to early repolarization, shaping the phase 1 notch of the action potential. In neurons, K_V_4.3 regulates action potential duration, firing frequency, and synaptic integration. To fully reproduce the *I*_tof_ and *I*_A_ currents, K_V_4.3 channels need to form complexes with several interacting proteins collectively known as K_V_4.3 channelosome. Among these, KChIPs are the most studied, and they enhance surface expression, delay inactivation kinetics, and accelerate both activation and recovery from inactivation kinetics. Disruption of K_V_4.3 function or its regulatory complex has been involved in a range of pathophysiological conditions, including cardiac arrhythmias, epilepsy, and neurodegenerative diseases.

Notably, small molecule ligands that target KChIPs have drawn considerable interest, as they significantly modify the biophysical properties of K_V_4.3/KChIP currents and represent promising tools for modifying channel function. Given the crucial role of KChIPs in regulating cellular electrophysiology, modulating their activity—either by enhancing or inhibiting their effect on K_V_4 channels—offers a promising strategy for developing therapies targeting K_V_4-related pathologies.

IQM-22110 was designed with the purpose to reach unexplored cavities on the known KChIP3 binding site. To this end, an additional aromatic ring at R_1_ position was incorporated on the (phenylacetamido)benzoic acid skeleton and docking studies highlighted that it may enhance compound affinity for KChIP3 due to additional hydrophobic interactions. Furthermore, IQM-22110 may stablish an additional interaction with Lys166, engaging a novel pocket. FRET studies based on the quenching of tryptophan fluorescence emission at 330 nm allowed to calculate the affinity of IQM-22110 to KChIP3 with an IC_50_ of 230 nM. As far as we are aware, this novel small molecule stands out as the most potent KChIP3 ligand identified.

In the present study we have also analyzed the electrophysiological effects of IQM-22110 on K_V_4.3 and K_V_4.3/KChIP3 channels, studying the concentration-, voltage- and use-dependent block of this new ligand on them. In this work, not only the concentration-dependent block of K_V_4.3/KChIP3 channels induced by IQM-22110 suggests the existence of two binding sites with two different IC_50_s (28 nM and 22 μM); but also, the different electrophysiological effects that IQM-22110 on K_V_4.3/KChIP3 current produced at concentrations in the nM and in the μM range. The main biophysical differences were detected in the voltage-dependent, the use-dependent block and in the kinetics of recovery from inactivation. At high concentrations, within the μM range, IQM-22110 binds to the K_V_4.3/KChIP3 channel in a similar way that it does to the K_V_4.3 channel alone. Moreover, under both experimental conditions, IQM-22110 (at μM range) accelerated the recovery from inactivation, inducing an overshoot, which resulted more prominent in K_V_4.3/KChIP3 channels due to their fast recovery from inactivation, suggesting that this ligand exhibits very low or null affinity from the inactivated state. This should explain why the IC_50_ of block induced by this compound on K_V_4.3 is lower than the IC_50(2)_ of block produced by IQM-22110 on K_V_4.3/KChIP3 channels, because the IQM-22110 access to this binding site in K_V_4.3 would be hindered by the presence of KChIP3. On the other hand, IQM-22110 at nM concentrations likely only binds to KChIP3 in a selective mode. Indeed, the FRET experiments do not show a biphasic dose-response curve of the binding of IQM-22110 to KChIP3 and the IC_50_ (230 nM) is in the range of the IC_50(1)_ (28 nM) obtained by the electrophysiology recording approach. These differences may be attributed precisely to the distinct techniques employed in both experiments, as well as to the presence of the K_V_4.3 channels in the electrophysiological experiments. Interestingly, these effects were not observed in K_V_4.3/KChIP2 channels. To the best of our knowledge, this is the first report of a KChIP3 ligand exhibiting this behavior, suggesting the existence of a high-affinity binding site in K_V_4.3/KChIP3.

Moreover, molecular dynamics studies highlighted Tyr130 and Lys166 as key residues for the IQM-22110 binding to KChIP3, with a distance to Trp169 of circa.10 Å, in agreement with the apparent distance tryptophan–ligand calculated by FRET experiments (circa. 10.02 ± 0.78 Å), while indicating that Tyr118 is not involved. Site directed mutagenesis studies conducted by electrophysiology showed that indeed both residues are critical for the binding of IQM-22110 to the high affinity site in K_V_4.3/KChIP3. This knowledge will guide the development of selective modulators of K_V_4.3/KChIP3 channels, laying the bases for drug discovery efforts targeting diseases in which these channels are implicated.

## 4. Conclusions

To the best of our knowledge, here we describe the first KChIP3 ligand that selectively modulates K_V_4.3/KChIP3 *versus* K_V_4.3/KChIP2 and K_V_4.3 channels. The high-affinity binding site of IQM-22110 to KChIP3 is primarily mediated by two critical residues, Tyr130 and Lys166. Collectively, these findings establish IQM-22110 as a valuable pharmacological tool and lay the groundwork for the development of novel modulators targeting K_V_4.3/KChIP3 channels in K_V_4 channelopathies.

## Methodology

### Synthesis of IQM-22110: 2-(2-([1,1’-biphenyl]-3-yl)acetamido)-4-chlorobenzoic acid (1)

2-([1,1’-biphenyl]-3-yl)acetic acid **2** (1.5 equiv) in SOCl_2_ (2 mL/mmol) was refluxed during 6h. After that, the solution was evaporated to dryness; the residue was dissolved in anhydrous THF (2 mL/mmol) and methyl 2-amino-4-chlorobenzoate (1.0 equiv) and propylene oxide (5.0 equiv) were added. The reaction was stirred overnight at room temperature, and, after that, the solvent was evaporated. The residue was dissolved in ethyl acetate (3 × 10 mL), washed with brine (30 mL), dried over Na_2_SO_4_. After purification by centrifugal circular chromatography using a gradient of 97% of hexane in ethyl acetate, the compound bearing a methyl ester was obtained with high purity. Next, a 2 N NaOH solution (0.22 mL) was added dropwise to a solution of the corresponding ester derivative (1 mmol) in THF/MeOH (1.33 mL/0.66 mL). After keeping the reaction mixture stirring at room temperature for 12 hours, the solvent was removed to dryness, and water (5 mL) was added. The solution was acidified to pH 3-4 using 1 N HCl. The aqueous phase was extracted with ethyl acetate (3 × 10 mL), and the combined organic layers were washed with brine (15 mL), dried over Na2SO4, evaporated to dryness and triturated with ethy. The residue was lyophilized from H2O/CH3CN (1:0.3) to yield the final compound **IQM-22110** (1) as amorphous solid and 97% of purity.

^1^H-NMR (400 MHz, DMSO-d_6_) δ (ppm): 13.83 (s, 1H), 11.32 (s, 1H), 8.65 (d, *J* = 2.2 Hz, 1H), 7.96 (d, *J* = 8.5 Hz, 1H), 7.71-7.64 (m, 3H), 7.59 (dt, *J* = 7.8, 1.4 Hz, 1H), 7.52 – 7.42 (m, 3H), 7.42-7.33 (m, 2H), 7.21 (dd, *J* = 8.5, 2.2 Hz, 1H), 3.88 (s, 2H). ^**13**^**C-NMR** (75 MHz, DMSO-d_6_) 8 (ppm): 170.0, 168.7, 141.8, 140.45, 140.0, 138.4, 135.2, 132.8, 129.8, 128.9, 128.6, 128.1, 127.5, 126.7, 125.4, 122.6, 119.1, 115.2, 44.6. **HPLC** (Sunfire C18, gradient from 50 to 95% of acetonitrile in water, 10 min) t_R_ = 5.96 min. LC-MS: 366.2 ([M+H]^+^)

### Induced Fit Docking (IFD) studies

The ligands were constructed using the Maestro interface of the Schrödinger suite of programs (Maestro, Schrödinger, LLC, New York, NY, 2021). The optimization of the ligand structures was carried out using the LigPrep module of the Schrödinger package with the OPLS_2005 force field [32]. This module corrects the distance and angles, creates the ionization states and performs an energy minimization. The proteins were prepared with the Protein Preparation tool within the Schrödinger suite of programs [33,34].

The molecular docking studies were carried out with the IFD protocol based on Glide and Prime, implemented in the Schrödinger package [35,36]. As the target protein different human KChIP3 homology models, previously build in the group on the base of NMR structures of *mus musculus* KChIP3 PDB code 2JUL, were used [37]. The binding site was selected as a cubic box of 10 Å centered on the residues Tyr118, Tyr130. The IFD protocol allows the flexibility of the residues within 5 Å from the ligand. First, a docking was performed with Glide reducing the Van der Waals radius of the amino acids of the binding site and the ligand by 50% [38–40]. Then, for each conformation obtained, the side chains of those amino acids within 5 Å from the ligand were optimized, using the Prime program (SCHRÖDINGER 2021a. Prime, Schrödinger, LLC, New York, NY, 2021) [41,42]. Those complexes within a 30 kcal/mol from the minimum were selected for redocking with the Glide program, using the default Glide parameters. The selection of the pose was based on IFDscore and on visual inspection. Visualization and creation of figures was performed with PyMOL Molecular Graphic System v2.5.2 (Schrödinger Inc., München, Germany). These simullations were carried out on the serves of *Centro Técnico de Informática* (CTI-CSIC).

### Molecular dynamics (MD) studies

The MD studies were carried out using Amber 20 [43] and the FF19SB force field for the protein and GAFF for the ligand. To perform MD, each protein-ligand complex was solvated in an octahedral box of water molecules using the TIP3P water model, with a maximum distance between the protein and the edge of the box of 10 Å. Next, the necessary counterions were added to neutralize the system. Periodic boundary conditions were used. First, a minimization of 30.000 steps was carried out, initially with a positional restraint weight of 5.0 Kcal mol^-1^ A^-2^, restraints were progressively weakened and removed in the last 10.000 steps. Subsequently, the system was equilibrated over 100 ps using a positional restraint of 5.0 Kcal mol^-1^ A^-2^ with integration time steps of 1.0 fs. The water was equilibrated for 75 ps, while the solutes were maintained restraint. The restrains were progressively weakened until they were removed in the last 25 ps. Once the system was equilibrated, a production phase of 1000 ns was carried out, maintaining constant the temperature and pressure (NTP ensemble). The temperature was maintained at 298 K using the Langevin thermostat. The time step was set to 2 fs, and the SHAKE algorithm was used. The electrostatic interactions were calculated using the PME method (particle mesh Ewald). The visualization and creation of figures were performed with PyMOL Molecular Graphic System v2.5.2 (Schrödinger Inc., München, Germany). These calculation were carried out on DRAGO (*Centro Técnico de Informática*, CTI-CSIC) and FinisTerrae III (*Centro de Supercomputación de Galicia*, CESGA).

### Fluorescence Binding Assay

Fluorescence binding assay was carried out on a Jasco FP-8500 spectrofluorometer (JASCO Deutschland GmbH, Pfungstadt, Germany). Different IQM-22110 concentrations were added to KChIP3 (0.5 µM) in 50 mM HEPES pH = 7.4, 100 mM NaCl and 1 mM CaCl_2_ at room temperature. Fluorescence emission was scanned from 280 nm to 650 nm (1 s integration time) with an excitation wavelength of 260 nm and slit widths of 5 nm for excitation and emission. All the spectra were corrected for background fluorescence by subtracting a blank scan of the solvent solution. Fluorescence resonance energy transfer (FRET) between KChIP3 and IQM-22110 was monitored by the decrease of the intrinsic tryptophan fluorescence emission at 330 nm and analyzed using the equation:

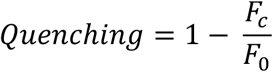

where *F*_*c*_ and *F*_0_ are the fluorescence emission at 330 nm in the presence and in the absence of IQM-22110, respectively. Normalized quenching at different IQM-22110 concentrations was analyzed by a Hill equation:

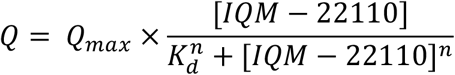

where Q_max_ represents the calculated maximal quenching, [IQM-22110] the IQM-22110 concentration, K_d_ the equilibrium dissociation constant and n is the Hill coefficient, using Origin 2023 software (Origin Labs Inc., Northampton, MA, USA). Three complete repeats were carried out and error bars are reported as the standard deviations (s.d.). We determined the distance (*R*) between the donor (Trp169) and IQM-22110 using the following formula:

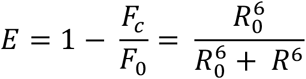

where the energy transfer efficiency from donor to acceptor (*E*) is related to the fluorescence intensity of the donor in the absence (*F*_*0*_) and the presence (*F*) of the acceptor at saturating concentration. The distance in which energy transfer is 50% (*R*_*0*_) was calculated using the equation:

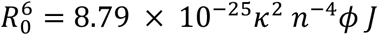

considering the orientation factor (*k*^*2*^) as 2/3, the quantum fluorescence yield (*f*) of tryptophan as 0.037 (cite: Gonzalez WG, Pham K, Miksovska J. Modulation of the voltage-gated potassium channel (K_V_4.3) and the auxiliary protein (KChIP3) interactions by the current activator NS5806. J Biol Chem. 2014 Nov 14;289(46):32201-32213. doi: 10.1074/jbc.M114.577528.), and the refractive index of the medium (*n*) as 1.4. The degree of spectral overlap (*J*) was estimated by the spectral overlap between donor and acceptors according to

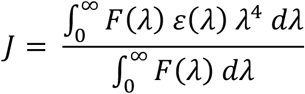

where *F(λ)* is the fluorescence of KChIP3, and *ε(λ)* is the molar absorptivity coefficient of IQM-22110.

### Cell culture and transfection

The experiments were performed in CHO-K1 cells from the American Type Culture Collection (Rockville, MD, USA). Cells were cultured in Iscove’s modified Eagle’s medium supplemented with 10% (v/v) fetal bovine serum (FBS), 1% (v/v) L-Glutamine and antibiotics (100 IU/ml penicillin and 100 mg/mL streptomycin), all from Gibco, Paisley, UK, at 37ºC in a 5% CO_2_ atmosphere. The culture medium was changed every 2-3 days and, after reaching confluence, the cultures were trypsinised with TrypLE™ Express (Life Technologies).

Cells were transiently transfected with Kv4.3 (cloned into pBK-CMV) with or without KChIP3 (AF199599.1, cloned into pcDNA3.1, provided by J.R. Naranjo (Centro Nacional de Biotecnologia, CSIC, Madrid, Spain)). In both cases, cells were cotransfected with EBO-pcDLeu2 as a reporter gene, codifying CD8. For Kv4.3/KChIP2 currents, 1 µg KChIP2 (cloned into IRES mCherry using *XbaI* and *XhoI* restriction sites) was cotransfected with Kv4.3. Transfection was performed using Fugene-6 (Promega) following manufacturer’s instructions as previously described [12,27,44]. After 48 h transfection, cells were removed from culture plates using TrypLE™ Express (Life Technologies), after exposing them to polystyrene microspheres bound to anti-CD8 (Dynabeads M-450, Thermo Fisher Scientific; [45,46]) for Kv4.3 and Kv4.3/KChIP3 currents, or selected with Fluorescence Activated Cell Sorting (FACS) for Kv4.3/KChIP2 currents (positive to mCherry) were isolated using procedure and preserved in DMEM until being used.

### Site-directed Mutagenesis

We used two mutations (KChIP3^Y118A^ and KChIP3^Y130A^) previously reported [37]. The point-mutation K166A in KChIP3 (c.496A>G, c.497A>C, p.K166A) was introduced KChIP3^WT^ by PCR, using Herculase II Fusion DNA polymerase (600675, Agilent, Santa Clara, CA USA) similarly to previously described [47]. The following primers were used (modified sequence is shown in bold red and lower-case letters):

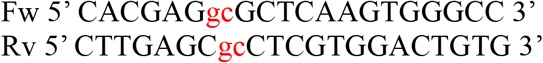

The PCR product was purified with QIAquick Gel Extraction Kit (Qiagen) and then digested with *DpnI* (New England BioLabs) for 1 h at 37°C and 15 min at 70°C in order to remove the traces of the original vector and obtain only the DNA resultant of the PCR. Wild-type and mutant constructs were verified by DNA sequencing.

### Whole-cell Patch-clamp

Electrophysiological experiments were performed using the whole-cell configuration of the patch-clamp technique. The intracellular pipette solution contained the following (in mM): K-Aspartate 80, KCl 42, phosphocreatine 3, KH_2_PO_4_ 10, Mg-ATP 3, HEPES-K 5, EGTA-K 5 (adjusted to pH=7.25 with KOH). The bath solution contained the following (in mM): NaCl 145, KCl 4, CaCl_2_ 1.8, MgCl_2_ 1, HEPES-Na 10 and glucose 10 (adjusted to pH=7.4 with NaOH). Currents were recorded using an Axopatch 200B amplifier and a Digidata 1440A analog-to-digital converter using the PCLAMP 10 (Molecular Devices) and data were stored on a personal computer (Hewlett Packard). Currents were recorded at room temperature (21–23°C) at a stimulation frequency of 0.1 Hz, low pass-filtered and sampled at 2 kHz. Borosilicate pipettes (GD1, Narishige) were pulled using a programmable patch micropipette puller (P-87, Sutter Instrument) and were heat polished with a microforge (MF-83, Narishige). The average pipette resistance ranged from 3 to 4 MΩ and GΩ (2–5 GΩ) seal formation was achieved by suction. OriginPro 2023b and the CLAMPFIT 10.6 utility of the PCLAMP 10 software were used to perform least-squares fitting and data presentation.

To record K_V_4.3, K_V_4.3/KChIP2, K_V_4.3/KChIP3 (wt or mutants) currents, the membrane potential was held at -80 mV and the interpulse interval was set to a minimum of 10 s. Current-voltage relationship (I-V) was obtained by the application of 250-ms pulses between –80 mV and +60 mV in 10 mV increments followed by pulses to –40 mV to record the deactivating tail currents. Voltage dependent of channel activation was analyzed by plotting the maximum current amplitude of the deactivating currents vs. the membrane potential of the previous pulse.

IQM-22110 perfusion was monitored with test pulses from -80 mV to +60 mV applied every 10 s until steady state block was obtained (within 3 to 5 min) by using the CLAMPEX

11.1 utility of PCLAMP 11 software (Molecular Devices Co., San Jose, CA, USA). Other pulse protocols are described in the Results Section.

Block produced by IQM-266 was calculated after measuring the current at the maximum peak or under the area of the current after applying different concentrations of the compound (0.001–150 μM). The percentage of block was calculated by:

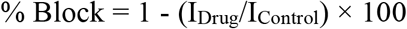

From the fitting of these values to a Hill equation, concentration-effect curves were generated, obtaining the values of the IC_50_ and the Hill coefficient (n_H_).

Activation and inactivation were fitted to a monoexponential process with an equation of the form:

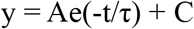

where τ represents the system time constant, A represents the amplitude of the exponential, and C is the baseline value. The voltage dependent of the activation and the inactivation curves were fitted to a Boltzmann equation:

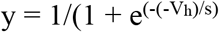

where s represents the slope factor, V represents the membrane potential, and V_h_ represents the voltage at which 50% of the channels are open or inactivated (activation or inactivation curves, respectively). In all cases, the control and the experimental condition was the same cell before and after being exposed to IQM-22110 (see Results Section).

### Statistical Analys

Data are expressed as mean ± SEM of n measurements. Comparisons between mean values were performed by using the Student’s t-test or ANOVA, as appropriate (Bonferoni’s correction in post–hoc comparisons). Statistical significance was defined as P < 0.05. Sample size (n) is reported in figure legends.

## Funding and Akcnowledgements

This work has been funded by grants: PID2022-137214OB-C21 (to CV) and PID2022-137214OB-C22 (to MG-R) funded by MCIN/AEI/10.13039/501100011033 and by “ERDF A way of making Europe”; PID2023-148243OB-I00 AEI/10.13039/501100011033 (to AO); Grant CB/11/00222 funded by Instituto de Salud Carlos III CIBERCV (to C.V.); CSIC grants 202580E109 (to MMM and MG-R) and 2023AEP049 (to CV); P. G. Socuéllamos was funded by FPU17/02731 granted by Spanish Minister of Science and Innovation. C. Viedma-Barba holds a FPI Predoctoral contracts subsidized by Spanish Minister of Science and Innovation (PRE2020-093542) granted by MCIN/AEI/10.13039/501100011033 and “ESF Investing in your future”. A. de Benito-Bueno holds a FPI grant BES-2017-080184. M. Redondo holds a FPI Contract subsidized by Spanish Minister of Science and Innovation (PREP2022-000706). A.P.-L. acknowledges funding by a Ramon y Cajal grant (RYC2018-023837-I) and Max Planck Society through the funding of a Max Planck Partner Group on “Regulation of the SNARE zippering by complexins and synaptotagmins” at the University of Granada.

We thank Centro de Supercomputación de Galicia (CESGA) for performing molecular dynamics studies using the Finis Terrae III supercomputer.

